# Age-linked heterogeneity among oligodendrocyte precursor cells in the cerebral cortex of mice and human

**DOI:** 10.1101/799544

**Authors:** A Bribián, EM Medina-Rodríguez, F Josa-Prado, I García-Álvarez, I Machín-Díaz, V Murcia-Belmonte, L Vega-Zelaya, J Pastor, L Garrido, F de Castro

## Abstract

Oligodendrocyte precursor cells (OPCs) are responsible for spontaneous remyelination after a demyelinating lesion. They are present in large parts of the mouse and human central nervous system, both during development and in adulthood, yet how their physiological behaviour is modified throughout life remains largely unknown. Moreover, the activity of adult human OPCs is still not fully understood. Significantly, most of the molecules involved in OPC-mediated remyelination are also involved in their development, a phenomenon that may be clinically relevant. In this article, we have systematically analyzed the intrinsic properties of OPCs isolated from the cerebral cortex of neonatal, postnatal and adult mice, as well as those recovered from neurosurgical adult human cerebral cortex tissue. We also analyze the response of these cells to two molecules that have known effects on OPC biology during development and that are overexpressed in individuals with Multiple Sclerosis (MS): FGF2 and anosmin-1. By analyzing intact OPCs for the first time with H-1 HR-MAS NMR spectroscopy, we show that these cells behave distinctly and that they have different metabolic patterns in function of their stage of maturity. Moreover, their response to FGF-2 and anosmin-1 differs in relation to their developmental stage and in function of the species. Our data reveal that the behaviour of adult human and mouse OPCs differs in a very dynamic way that should be considered when testing drugs and for the proper design of effective pharmacological and/or cell therapies for MS.

## INTRODUCTION

Oligodendrocyte precursor cells (OPCs) are the sole source of oligodendrocytes, the myelin-forming cells in the central nervous system (CNS), and they are widely distributed throughout both the gray and white matter [1-5]. The vast majority of mature oligodendrocytes are born during postnatal life but in adults, oligodendrocytes can also be generated from OPCs that continue to divide and generate new-myelinating cells during adulthood [5-11]. In the adult brain, OPCs constitute 2–3% of the cells inthe gray matter and 5–8% in the white matter [1]. These cells are maintained as a resident population by self-renewal, representing the main proliferative cell type outside the neurogenic regions of the CNS [1,7,12,13]. However, the adult OPC population is not homogeneous as OPCs cycle less intensely with age, their proliferation rate is higher in the whiter matter than in the grey matter, and their differentiation rate declines progressively in postnatal and adult life [1,5,8-11,14,15]. Although the functions of adult-born oligodendrocytes remain largely unclear, they apparently maintain physiological oligodendroglial homeostasis and under pathological circumstances, they mediate long-term repair in the white matter after injury and disease, even in MS where they can partially restore the lost myelin sheaths [16,17]. In demyelinating diseases, there is an up-regulation of different molecules in the microenvironment in and around the lesions, including several factors that are important for developmental oligodendrogliogenesis, such as secreted semaphorins, growth factors, mitogens and adhesion molecules [18-24].

In the present study, we focused on cerebral cortex OPCs to avoid any variability introduced by anatomical origin or location. We examined the degree of intrinsic heterogeneity (both heterochronic and interspecies) of these cortical OPCs and we described the dynamic changes in their intrinsic properties when assessed by NMR (i.e, metabolic patterns), and how these changes affect their response to two molecules known to influence OPC biology during development and remyelination: FGF2 and anosmin-1 [19,25-33]. We systematically studied mouse cortical OPCs at different ages and OPCs isolated from the human adult cerebral cortex obtained from neurosurgery biopsies (resection margins of tumors and/or epilepsy surgery [34]). In terms of the latter, the scarcity of articles dealing with human OPCs is especially remarkable [21,34-38].

The data we obtained highlight the intrinsic physiological differences between mouse OPCs isolated at different ages. Indeed, while OPC proliferation and migration were enhanced similarly by FGF2, this factor produced different effects on OPC differentiation. By contrast, anosmin-1 elicited diverse effects on differentiating, and especially on migrating OPCs, yet not on the proliferation of these cells. While FGF2 was a chemoattractant for human and murine OPCs, anosmin-1 only attracted tumor-related cells and it repelled non-tumor OPCs. Neither FGF2 nor anosmin-1 affected human adult OPC differentiation. These observations confirm the chronological and inter-species heterogeneity of OPCs, adversity that will probably be relevant when designing effective pharmacological and/or cell therapies to repair myelin loss in MS and other demyelinating diseases.

## MATERIALS AND METHODS

### Animals

CD-1 mice were obtained from (Charles River Laboratories, Inc., Wilmington, MA) and maintained in the animal facilities of the Hospital Nacional de Parapléjicos (Toledo, Spain). OPCs were obtained from different age animals: embryonic (E16); postnatal (P0, P7, P15); and adult (P60). All experimental procedures were carried out in accordance with Spanish (RD223/88) and European (2010/63/EU) guidelines, and were approved by the Animal Review Board at the Hospital Nacional de Parapléjicos (SAPA001).

### Human biopsies

Biopsies of human tumor (resection of the safety margins but not the tumor itself) and non-tumor tissue (epilepsy surgery) were obtained from the Neurosurgery Service at the Hospital La Princesa (Madrid, Spain). All human samples were obtained from the adult brain cortex and the biopsies were predominantly from the temporal lobe (Table 1). They were transported at 4°C as rapidly as possible in AQIX® RS-I (AQIX Ltd., Imperial College BioIncubator) to reduce cell damage. All experiments involving human samples were carried out in accordance with the guidelines of the Research Ethics Committee of our institution, which approved our research, and all the subjects provided their written informed consent.

**Table 1:**
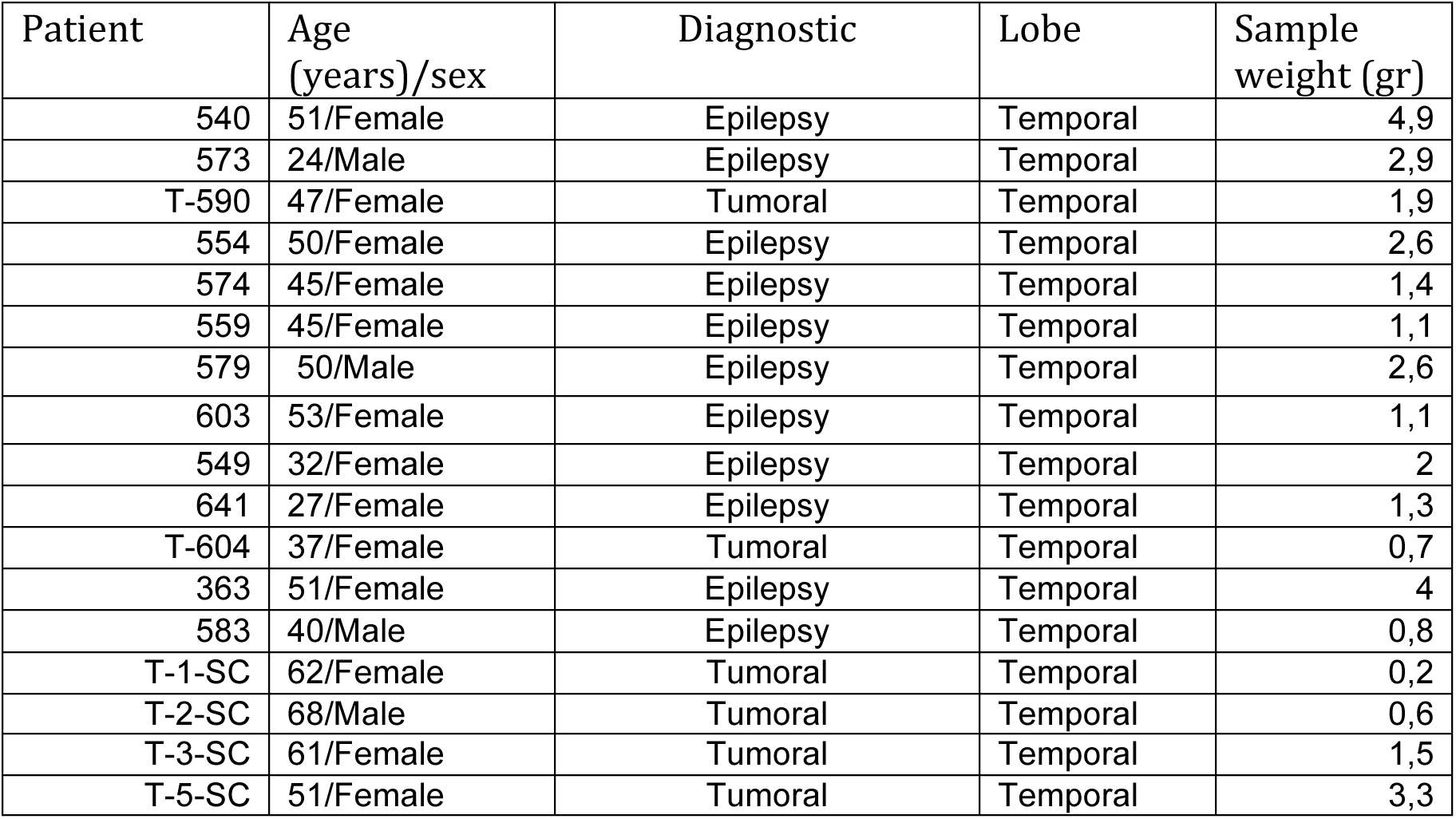
Summary of human samples used for in vitro experiments

| Patient | Age<br>(years)/sex | Diagnostic | Lobe | Sample<br>weight (gr) |
| --- | --- | --- | --- | --- |
| 540 | 51/Female | Epilepsy | Temporal | 4,9 |
| 573 | 24/Male | Epilepsy | Temporal | 2,9 |
| T-590 | 47/Female | Tumoral | Temporal | 1,9 |
| 554 | 50/Female | Epilepsy | Temporal | 2,6 |
| 574 | 45/Female | Epilepsy | Temporal | 1,4 |
| 559 | 45/Female | Epilepsy | Temporal | 1,1 |
| 579 | 50/Male | Epilepsy | Temporal | 2,6 |
| 603 | 53/Female | Epilepsy | Temporal | 1,1 |
| 549 | 32/Female | Epilepsy | Temporal | 2 |
| 641 | 27/Female | Epilepsy | Temporal | 1,3 |
| T-604 | 37/Female | Tumoral | Temporal | 0,7 |
| 363 | 51/Female | Epilepsy | Temporal | 4 |
| 583 | 40/Male | Epilepsy | Temporal | 0,8 |
| T-1-SC | 62/Female | Tumoral | Temporal | 0,2 |
| T-2-SC | 68/Male | Tumoral | Temporal | 0,6 |
| T-3-SC | 61/Female | Tumoral | Temporal | 1,5 |
| T-5-SC | 51/Female | Tumoral | Temporal | 3,3 |

### HOG cells

An immortalized oligodendroglioma cell line, HOG, was grown at 37°C in an atmosphere of 5% CO_2_, and in Dulbecco’s Modified Eagle’s Medium (DMEM, Gibco) supplemented with 10% Fetal Bovine Serum (FBS, Gibco), 100 U/mL penicillin and 100 µg/mL streptomycin (Sigma).

### Oligodendrocyte culture

OPCs were isolated as described previously [19,34,38]. Briefly, the cerebral cortex of mice aged E16, P0, P7, P15 and P60 was dissected out and the meninges were carefully removed. The cortical tissue was mechanically disaggregated and it was then enzymatically dissociated in a papain solution, filtered using a 100 µm nylon mesh strainer (BD Biosciences), and seeded in polyornithine-treated flasks in DMEM containing 10% FBS (BioWhittaker) and an antibiotic anti-mycotic solution (100 U/mL penicillin/0.1 mg/mL streptomycin and 0.25 µg/ml Amphotericin B; Sigma). The cultures were maintained at 37°C and in 5%CO_2_, and the medium was changed every 4 days, adding10 ng/ml of human PDGF-AA (Millipore) to the cultures from adult mice. When the cultures reached confluence, they were shaken overnight at 250 rpm to detach the oligodendrocyte progenitors growing on the top of the confluent astrocyte monolayer. The medium was then filtered through a 40µm nylon mesh strainer (BD Biosciences) and centrifuged at 900 rpm. The cells recovered were seeded twice (45 minutes each) in bacterial grade Petri dishes (Sterilin) to remove the adherent microglia and after another round of centrifugation, the resulting pellet was resuspended and the OPCs were counted and seeded.

For cells obtained from human biopsies, the same protocol was used with certain modifications published previously [34]. This protocol allowed us to obtain sufficient human OPCs to perform the necessary assays. Briefly, rather than 75 cm^2^flasks, 25 cm^2^ flasks were used with a final volume of 5 ml rather than 10 ml OPC medium per flask. In addition, the medium was supplemented with 10 ng/ml of human PDGF-AA (Millipore), as in the cultures obtained from adult mice. Cultures reached confluence after 30-40 days and the flasks were shaken at 230 rpm instead of 250 rpm.

### Preparation of extracts for anosmin-1

Proteins from the extracellular matrix (ECM) of CHO cells (CT) and CHO cells stably transfected with an expression plasmid carrying a C-terminal hemaglutinin-tagged (HA) version of the human KAL1 cDNA (A1 [39]) were extracted as described previously [32]. Briefly, the cells were washed once with calcium/magnesium-free Hank’s Balanced Salt Solution (Gibco) and they were then incubated at 4°C for 30 minutes in a gently rocking 10 cm diameter culture dish in 1 ml of 20 mM phosphate buffer (PB, pH 7.4), containing 350 mM NaCl and complete EDTA free protease inhibitor (Roche). The ECM proteins released into the buffer were concentrated ten-fold with Amicon Ultra-4 Ultracel-30k (Millipore Corporation) and used in the chemotaxis experiments (ECM extracts). The presence of anosmin-1 was confirmed in western blots probed with a peroxidase-conjugated rat monoclonal antibody against HA (High Affinity 3F10; Roche). Equal amounts of protein from untransfected and transfected cells were used in all the *in vitro* experiments described below.

### Chemotaxis

To test chemoattraction, cells (murine and human OPCs, and HOG cells) were seeded in the upper chamber of a chemotaxis chamber (40,000 cells/transwell), and FGF2 (20ng/ml; R&D Systems) or 0.1 mg/ml of CT and A1 cell extracts (see below) were added to the lower chamber in Bottenstein-Sato (BS) medium, as reported previously [19,27,32,40]. After 20 hours at 37°C, the cells were fixed in 4% paraformaldehyde (PFA) for 10 minutes at room temperature and the OPCs were stained with antibodies against Ganglioside GT3 (A2B5, 1:20; Developmental Studies Hybridoma Bank) and the Olig-2 oligodendroglial transcription factor (1:200, Millipore), and HOG cells were stained with Hoechst. After immunostaining, the filters were examined using an In Cell Analyzer 1000 (GE-HealthCare) and 15-20 microphotographs from each membrane were taken randomly. The In Cell Analyzer 1000 Work station software was used (GE-HealthCare) to quantify the number of transmigrated OPCs per field, expressing the data as the percentage of migrating OPCs relative to the controls±s.e.m. (Considered as 100% [32]). The data were analyzed with the Sigma Stat software package (SPSS).

### Time-Lapse Imaging

For a detailed study of OPC migration, time-lapse video-microscopy (Leica, DMI 6000B) was performed to observe the behaviour of the cortical OPCs. Purified OPCs were placed on poly-L-lysine and laminin coated coverslips (10,000 cells/coverslip; Sigma) and FGF2 (20ng/ml; R&D Systems) or CT CHO or A1-expressing CHO cell extracts (0.1 mg/ml, see above) were added in BS medium. Images were taken every 20 minutes for 18 hours and the speed of cell motility was assessed with the manual tracking plugin of the ImageJ software. The values represent the percentage of the mean velocity relative to the respective controls (± s.e.m.).

### Differentiation assay

Purified OPCs were placed on coverslips coated with poly-L-lysine and laminin (20,000 cells/coverslip), and FGF2 (20ng/ml; R&D Systems) or total protein from CT CHO cells or A1-expressing CHO cells (0.1 mg/ml, see above) were added in differentiation medium (BME:F12 media [1:1] supplemented with 100 µg/ml transferrin, 20 µg/ml putrescine, 12.8 ng/ml progesterone, 10.4 ng/ml selenium, 25 µg/ml insulin, 0.8 µg/ml thyroxine, 0.6% glucose, and 6.6 mM glutamine, as reported previously [34,38,41]. AraC (5µM; Sigma-Aldrich) was added to the cultures to inhibit proliferation and after 5-7days *in vitro* (DIV), the cells were fixed for immunocytochemical detection of CNPase (1:200; Covance) and Olig-2 (1:200; Millipore). Finally, 10 random fields per coverslip were photographed under a Leica microscope using a20X objective and the proportion of CNPase^+^ cells was assessed relative to the respective controls (± s.e.m.).

### Pharmacological treatments

In order to study the mechanism underlying the activity of the FGF2/FGFR1/anosmin-1 system in specific aspects of cortical OPC biology (chemoattraction, motogenicity, proliferation and differentiation), these processes were analyzed in the presence of the specific FGFR inhibitor SU5402 [19,27,32,40]. The SU5402 inhibitor was reconstituted in DMSO and the controls were administered the vehicle alone at the same concentration to account for any negative effects of DMSO on OPCs.

### Differential FGFR expression

FGFR1 expression in murine and human OPCs was assessed by dual immunofluorescence with the OPC marker NG2 (1:200; Millipore) and an antibody against FGFR1 (1:200; Santa Cruz Biotechnology). In order to understand the response to FGF2 and anosmin-1 observed in OPCs from different ages, we studied the expression of different FGFRs: FGFR1, FGFR2 and FGFR3. Cells were plated in 96-well tissue culture plates (10,000 cells/well) and maintained at 37°C in a 5% CO_2_ atmosphere and with 95% relative humidity in BS medium. After 1 DIV, the cells were fixed for 10 min at room temperature with 4% PFA. The OPCs were immunostained with antibodies against FGFR1 (1:200; Santa Cruz Biotechnology), FGFR2 (1:50; Santa Cruz Biotechnology) and FGFR3 (1:20; Santa Cruz Biotechnology), and against tubulin (1:40,000; Sigma-Aldrich). Antibody binding was detected using 680 or 800 IRDye conjugated secondary antibodies and the plates were scanned using the Odyssey Infrared Imaging System (LICOR). Fluorescence intensity was measured according to the manufacturer’s instructions and the amount of FGFRs relative to the amount of tubulin was normalized to the FGFR expression in P0 samples. The values represent the mean (± s.e.m.).

### Proliferation assays

Purified OPCs were seeded on poly-L-lysine and laminin coated coverslips (20,000 cells/coverslip), and FGF2 (20ng/ml; R&D Systems) or CT CHO or A1-expressing CHO cell extracts (0.1 mg/ml, see above) were added in BS medium. After 42 hours in culture, a BrdU pulse (6 hours) was administered and 24 hours later, the cells were fixed for immunocytochemical detection of BrdU and Olig-2, as described previously [27]. Finally, 10 random fields per coverslip were photographed under a Leica microscope using a 20X objective and the data were expressed as the percentage of proliferating OPCs relative to the controls (±s.e.m.) considered as 100%.

### H-1 HR-MAS NMR spectroscopy of OPCs

H-1 HR-MAS NMR measurements were performed on a Bruker spectrometer operating at 9.4 T (proton Larmor frequency of 400.14 MHz). The spectra were acquired at 4 °C to minimize the effect of temperature on cell stability during the acquisition time and to reduce the variations in some amino acids [42]. Sample spinning at the magic angle was applied at a speed of 2.4 kHz. The acquisition sequence and parameters were selected to assess the presence of metabolites of interest and their mobility, essentially using T2-filtered (CPMG) 1D experiments recorded with relaxation times of 2 and 60 ms. Low power presaturation during the interscan delay of 3 s was applied for water suppression. Typically, 256 scans were accumulated using a spectral width of 20 ppm, and the total acquisition time per spectrum was 13 min.

The spectra were processed using MestReNova version 8.1 software (Mestrelab Research, Santiago de Compostela, Spain) and all free induction decays were processed with exponential multiplication (0.5 Hz line-broadening) prior to Fourier transformation, followed by baseline correction. The chemical shifts were referenced as follows: a small amount (5 mL) of 10 mM DSS in D_2_O was added to one sample of intact cells and referenced to DSS = 0 ppm, after which all the spectra were processed identically and aligned (using the creatine methyl resonance at 3.026 ppm) with respect to the sample with added DSS. Of the various methods to estimate or measure the intensity of NMR signals [43], we compared peak intensity ratios using the methyl signal of creatine as an internal reference. Creatine is commonly used as an internal concentration reference for in vivo H-1 NMR because its concentration correlates with the number of metabolically active cells and it can be used as a measure of viable cell number [44]. Cells were washed (three times) with a deuterated PB solution, centrifuged, and the cell pellet was frozen in liquid nitrogen and stored at −80 °C. Typically, approximately 60 µL of the cell pellet was placed into a 4 mm Ø zirconia rotor for each sample.

### Imaging analysis

Fluorescence digital images were obtained with a DFC480 FX digital camera (Leica) coupled to a Leica DM5000 B microscope or using a SP5 resonant scanner (Leica Microsystems).

### Statistical Analysis

For all experimental conditions, the results were analyzed using a*t*-test (SigmaPlot 11 software (SPSS Science, Inc.) and critical values of: *P<0.05, **P<0.01 and ***P<0.001.

## RESULTS

### Changes in the intrinsic properties of OPCs with aging

The intrinsic properties and metabolic pathways of murine OPCs isolated from the cerebral cortex were assessed at different ages (P0, P15 and P60). For the first time, we assessed these changes using H-1 high-resolution (HR)-magic angle spinning (MAS) NMR spectroscopy, which allows a large number of metabolites or small molecules in the cytoplasm to be systematically evaluated and their interaction with the local environment probed [45-47]. This approach may therefore provide valuable information on cellular physico-chemical characteristics that may be correlated with cellular processes of interest. We obtained 3 spectra for each group of OPCs, with two different echo-times TE (“T2 filters T2”: 2, 10 and 60 ms), analyzing differences in signal intensities of peaks associated to the metabolites present in intact OPCs (see Table 2, Fig.1a). If we compared the proton spectra corresponding to P0, P15 and P60, for a given T_2_-filter (TE), we could see that there were not large differences between ages (Fig.1b). However, at TE of 60 ms, a relative increase in most peak intensities at P15 is observed (Fig 1c-d). At long echo times (TE: 60 ms), the intensity of the peaks associated to macromolecules or molecules with low mobility decreases (i.e., molecules bound to membranes or intracellular structures). It is remarkable that one the most significant (p < 0.05) signal change corresponds to myo-inositol, a metabolite known to be present in cell cytoplasm. Signal intensities depend on their T_2_ relaxation time, which is directly related to the molecular mobility in the cytoplasm. The higher metabolite mobility at P15 suggests a lower viscosity in the cytoplasm and low interaction between metabolites and the cytoskeleton, which as shown later, are associated with a decreased in the migratory capability and a decrease proliferation rate. These results indicate that OPCs isolated at P15 display intrinsic differences to those isolated at P0 or P60.

**Table 2.** Chemical shifts of main metabolites observed in OPCs

| Metabolite | $\delta$ (ppm), multiplicity, number H |
| --- | --- |
| myo-inositol | 3.93, m |
| myo-inositol | 3.77, m |
| glycine | 3.54, s, 2H |
| taurine | 3.42, t, 2H |
| hipotaurine | 3.30, t, 2H |
| total-choline | 3.23, s, 9H |
| glutamine | 2.44, m, 2H |
| glutamate | 2.34, m, 2H |
| glutamine/glutamate/NAA | 2.12 |
| acetate | 1.91, s, 3H |
| lipids | 1.70 |
| alanine | 1.47, d, 3H |
| lactate and lipids (methylene) | 1.31 |
| lipids and aminoacids (methyl) | 0.94, t, 3H |

**Figure 1:**
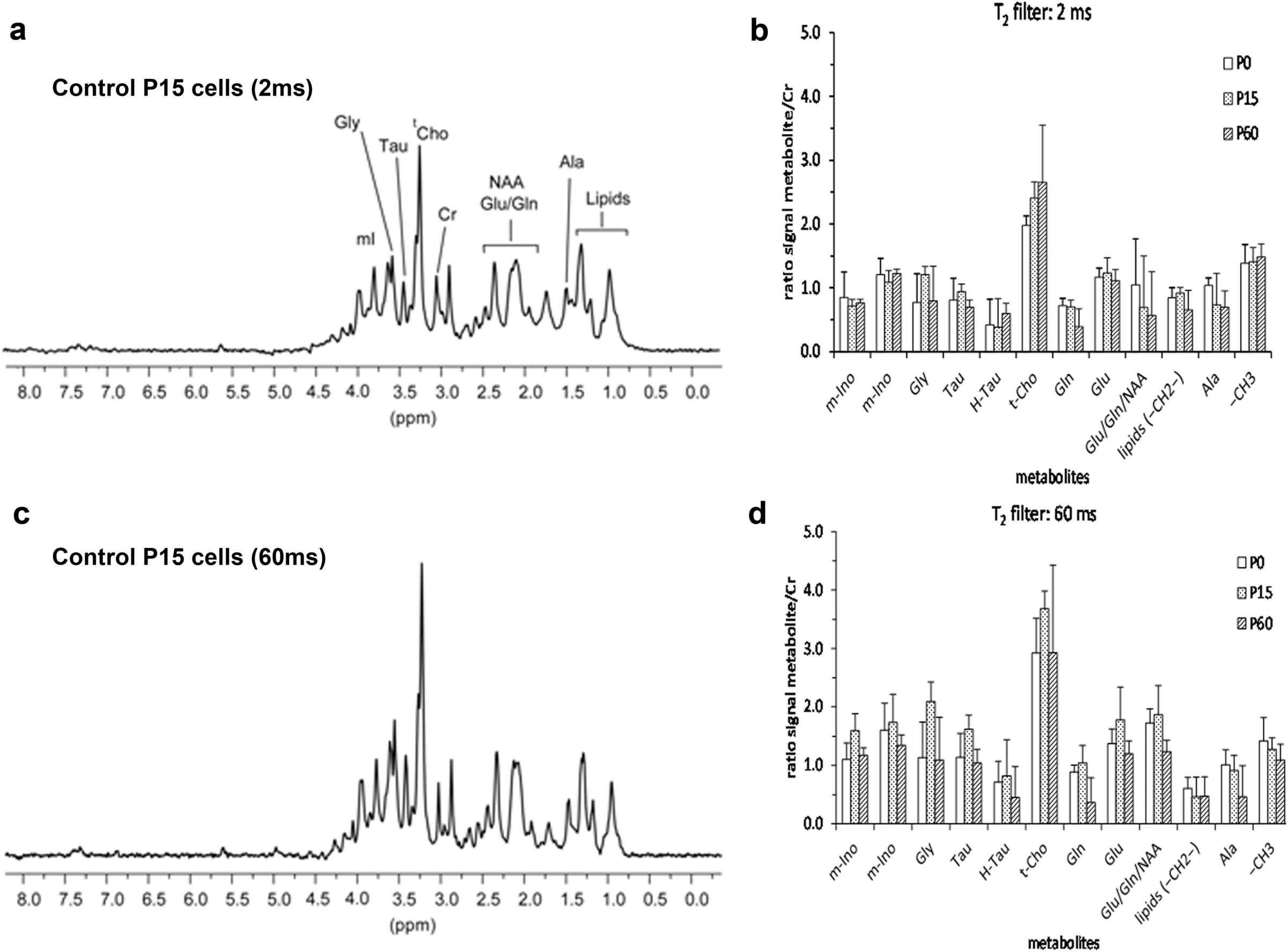
NMR of intact OPCs. Proton HR-MAS NMR spectra with water suppression corresponding to a sample of OPCs (P15) illustrating the chemical shift assignment of selected metabolites with T_2_-filters of 2 ms (a) and 60 ms (c), and the intensities of the metabolites indicated normalized with respect to creatine for T_2_-filters of 2 ms (b) and 60 ms (d). An increase in the relative intensities of P15 metabolites with respect to those of P0 and P60, for the T2-filter of 60 ms, is observed suggesting a decrease in cytoplasm viscosity.

Since the differences in the metabolic patterns may give rise to differences in other aspects of OPC behaviour, such as proliferation and migration, it was not surprising that the rate of proliferation was significantly higher at P0 and lowest at P15 (Fig. 2a). Notably, OPCs isolated from adults (P60) proliferated at an intermediate rate (Fig. 2a) and in general, there was a significant decrease in the migratory capacities of OPCs with age (Fig. 2b). This was corroborated by video time-lapse analysis, where P15 (the slowest) and P60 OPCs migrated slower than perinatal OPCs (Fig. 2c). Moreover, P60 OPCs stopped least often during their migration (Table 2) and in total; P0 OPCs migrated further while P15 OPCs migrated over the shortest distance (P0, 332µm; P15, 128µm; P60, 210 µm). These data highlight the heterogeneity in OPC behaviour (i.e., proliferation rate and migratory properties) under basal conditions, which could be related to differences in their metabolic pattern and variation in relaxation times of metabolites (i.e., cytoplasm viscosity). The strongest differences were detected in P15 OPCs, which may reflect the large scale myelination that is ongoing at that time [48,49].

**Figure 2:**
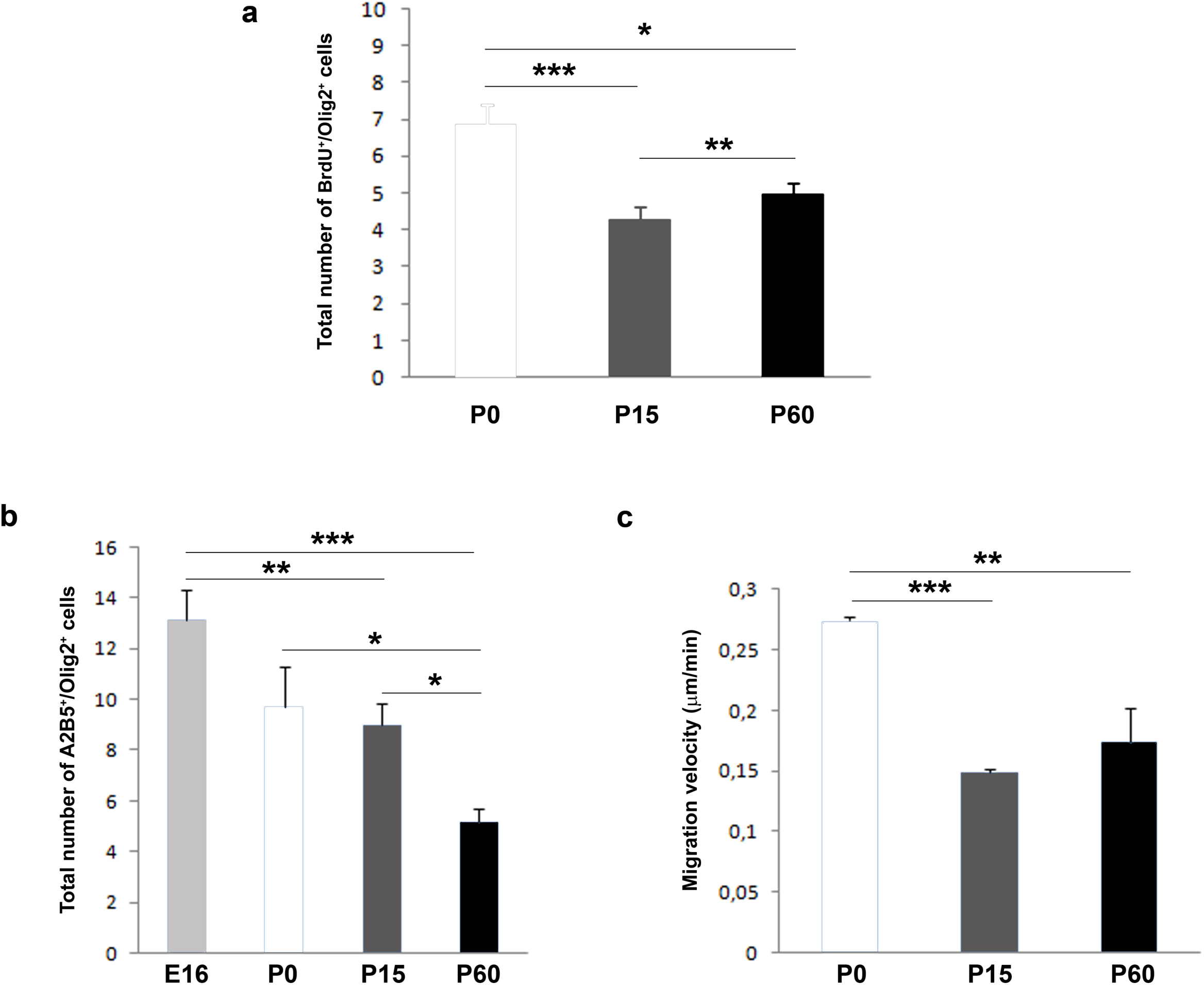
OPC proliferation and migration rates differ with age. a) Proliferation rate of neonatal, postnatal and adult OPCs. The number of proliferating OPCs also decreases with age. b) Transmigrated cells of different ages after 20 hours in control conditions: E16, P0, P15 and P60. The number of transmigrated cells decreases with age. c) Histogram represents the migration speed at P0, P15 and P60. The migration speed is higher at P0, and it diminishes in OPCs from P15 and P60 mice. For all experimental groups, the results were analyzed using ANOVA and a Student’s*t*-test: *P<0.05, **P<0.01, and ***P<0.001.

### Migrating OPCs respond dynamically to FGF2 and anosmin-1 in an age-dependent manner

FGF-2 and anosmin-1 exert distinct chemotropic effects on rat SVZ neuroblasts, and on rodent embryonic and postnatal OPCs, mainly acting via FGFR1 [19,27,32,33,40,50]. Hence, these molecules represent a good tool to systematically study the physiological heterogeneity in OPC populations, avoiding the variability in structure and origin by restricting our study to cerebral cortex OPCs. Exposure of OPCs isolated at different stages (E16, P0, P7, P15 and P60) to FGF2 homogeneously increased the number of transmigrating cells, confirming the previously described chemoattractive effect of this factor (and its potential motogenic effect as well: Fig. 3a). The effect of FGF2 on migration was significantly stronger on OPCs isolated from E16 and early postnatal stages (P0 and P7) than on those from later stages, reflecting a gradual loss of sensitivity with age and CNS maturation. Interestingly, anosmin-1 induced different OPC responses depending on the age of the cells: neutral for E16 OPCs; significant chemoattraction of OPCs isolated at P0 and P7; chemorepulsion in young adults (P15 and P60: Fig.3a). At all ages except P0, FGF2 triggered a stronger OPC response than anosmin-1 (Fig. 3a).

**Figure 3:**
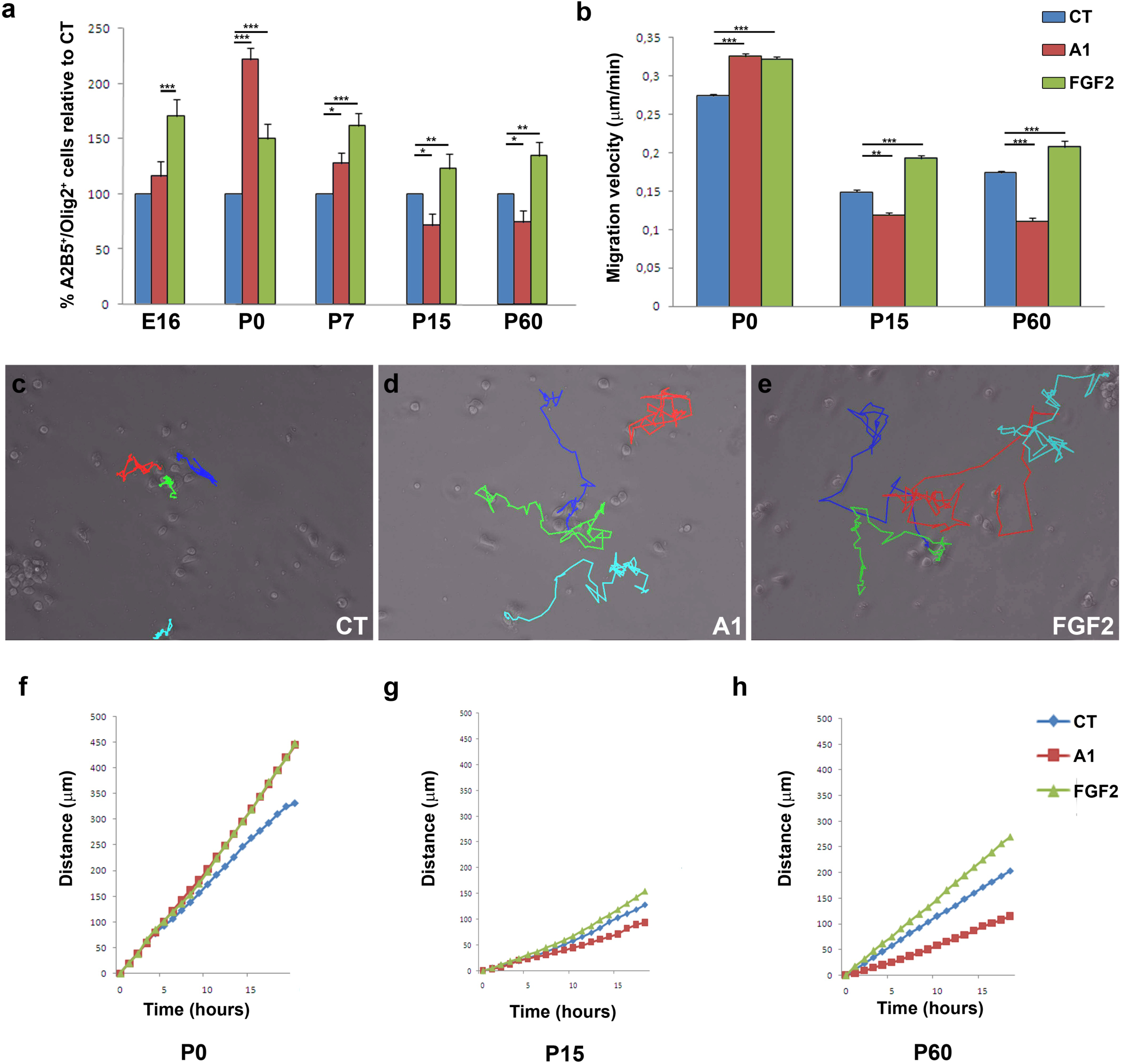
FGF2 and anosmin-1 control OPC migration. a) Histograms showing the proportion of OPCs relative to the controls quantified in the presence or absence of FGF2 or anosmin-1 at different stages: E16, P0, P7, P15 and P60. ANOVA showed anosmin-1 had no effect on migration at E16 and P7, in contrast to other stages, while and effect of FGF2 was evident at all stages. b) Histogram from the video time lapse analysis representing the speed of migration at P0, P15 and P60 in the presence of either factor. c-e) Images show the manual tracking at the end of the video time lapse recording for P0 OPCs, in control conditions (c) and in the presence of anosmin-1 (d) or FGF-2 (e). f-h) Analysis of the total distance migrated after 18hoursin all conditions: P0 (f), P15 (g) and P60 (h). All experimental groups were analyzed with a Student’s *t*-test: *P<0.05, **P<0.01, and ***P<0.001.

To more clearly distinguish between the chemotropic and motogenic effects of FGF2 and anosmin-1 (direction or rate of migration), we followed OPCs from different aged mice by video time-lapse microscopy. FGF2 increased the motility of OPCs at all the stages studied and it increased the velocity of OPCs by around 20% relative to their counterparts in control conditions (Fig. 3b). By contrast, anosmin-1 augmented the motility of P0 OPCs, yet it clearly reduced the speed of P15 and P60 OPCs (Fig.3b). Interestingly, P0 OPCs migrated further in the presence of either of the molecules tested than in control conditions, and they stopped less frequently (Table 2). On the other hand, P15 and P60 OPCs behaved similarly, migrating further and with fewer stops in the presence of FGF2, and anosmin-1 exerted exactly the opposite effects (Fig. 3c-h; Table 2). Together, our results suggest that both FGF2 and anosmin-1 mainly influence the motility of OPCs (motogenic effects), consistent with previous data from our group [19,27].

### Migration of human OPCs in the presence of FGF2 and anosmin-1

As indicated above, molecules involved in the migration of OPCs during development are up-regulated in the adult in conjunction with demyelination (including that occurring in MS lesions in humans), suggesting the need to test the effects of these molecules in adult OPCs. To complement this analysis, we sought to obtain information from human adult OPCs, evaluating the effects of both anosmin-1 and FGF2on three kinds of human OPCs in chemotaxis chambers: an oligodendroglioma cell line (HOG cells), and on OPCs isolated from tumor tissueand non-tumor tissue obtained through neurosurgery (the exact origin of these samples is indicated in the Methods). We first confirmed that these three cell types expressed FGFR1, the receptor thought to mediate the effects of FGF2 and anosmin-1 on OPC migration (Fig. 4a-f; [27,32,51-53]). It was not surprising to find that the tumor-derived OPCs were more motile in control conditions than non-tumoral OPCs (Table 3). However, exposure to the two factors tested also produced differences, whereby the number of transmigrated non-tumor OPCs significantly increased in the presence of FGF2 and was significantly lower in the presence of anosmin-1(Fig. 4g, j-l), response similar to those of P60 mouse OPCs (Fig. 3a). In addition, both proteins were motogenic and acted as chemoattractants for tumor-derived OPCs and HOG cells (Fig. 4h-l), as also reported for P0 murine OPCs (Fig. 3a). It should be noted that the effect of anosmin-1 was stronger on OPCs isolated from tumor samples than on the immortalized cell line. Video time-lapse analysis of non-tumor isolated OPCs revealed that anosmin-1 reduced their migration velocity and the distance they migrated, while increasing the number of stops, the opposite effect to that observed with FGF2 for the same three parameters (Fig.5a-j, Table 3), reflecting the contribution of pathological environment to the heterogeneity of OPC behaviour. Together, these results from adult OPCs demonstrated that they are a heterogeneous population of cells, in which the physiological heterogeneity depends on the species and age. Indeed, this highlights the risk of directly extrapolating from drug testing on OPCs isolated from neonatal/early postnatal rodent brains to the human adult scenario.

**Table 3:** Analysis of OPCs migration. The table shows the number of stops of migrating OPCs expressed as: number of stops/number of analyzed cells, in control conditions and in the presence of FGF2 and anosmin-1.

| Specie and age | CONDITION |  |  |
| --- | --- | --- | --- |
|  | CT | FGF2 | Anosmin-1 |
| Mouse P0 | 6.2 | 3.5 | 0.85 |
| Mouse P15 | 11.4 | 1.52 | 20.7 |
| Mouse P60 | 3.76 | 2.63 | 8.42 |
| Non tumoral human, adult | 3.61 | 8.8 | 17.6 |
| Tumoral human, adult | 23.7 | 19.7 | 23.1 |

**Figure 4:**
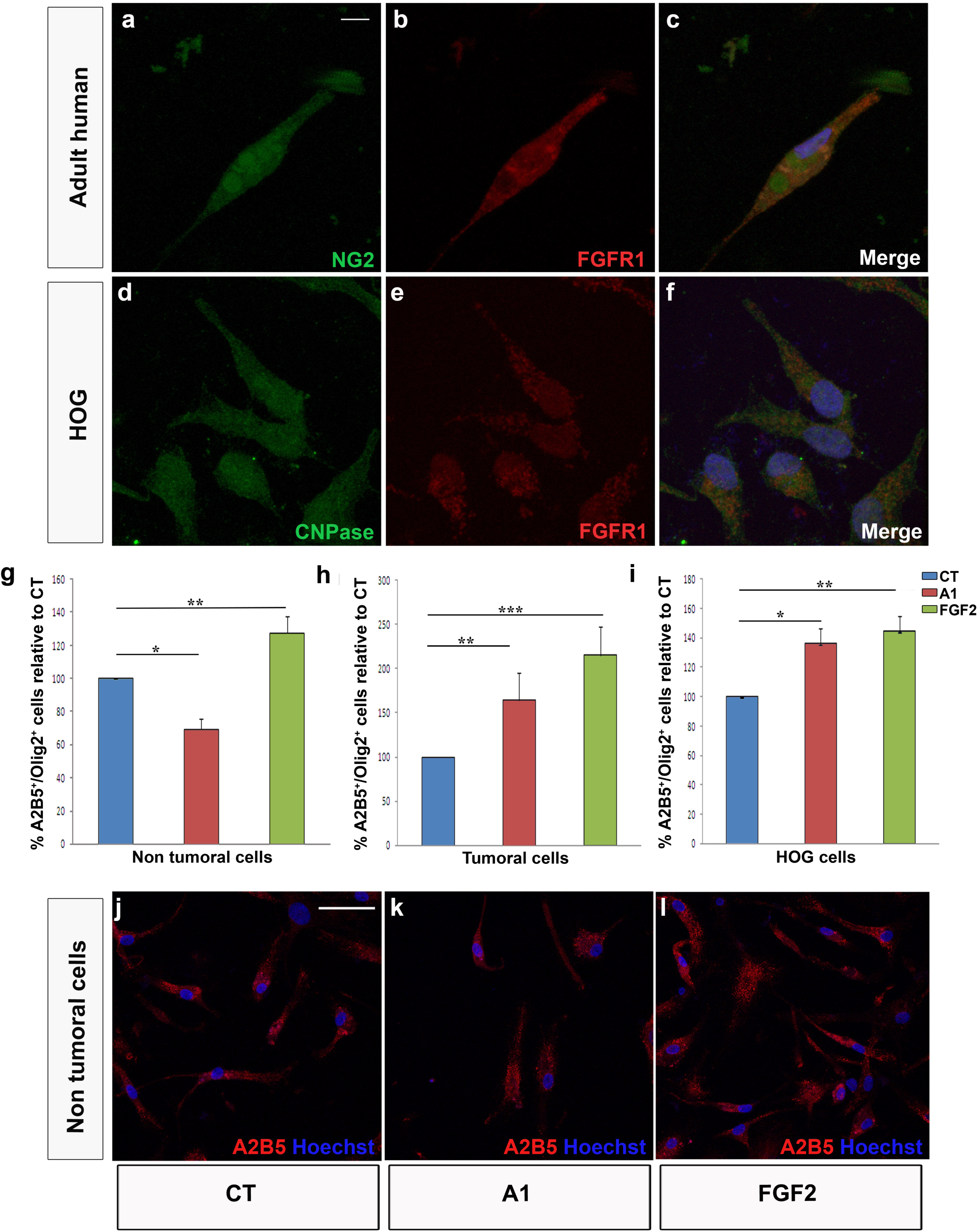
Effect of FGF2 and anosmin-1 on human OPC migration. Adult human OPCs (a-c) and HOG cells (d-f) expressed FGFR1. In both types of tumor cell analyzed, tumor cells (g) and HOG cells (h), the response to FGF2 and anosmin-1 was the same. Non-tumor cells responded to FGF2 like tumor cells, while the response to anosmn-1 is the opposite in these cell types (i). j-l) Images of tumor OPCs in chemotaxis chambers stained with A2B5 (red). Scale bar represents 7.5µm for a-f and 25 µm for j-l. The results were analyzed with a Student’s *t*-test: *P<0.05, **P<0.01, and ***P<0.001.

**Figure 5:**
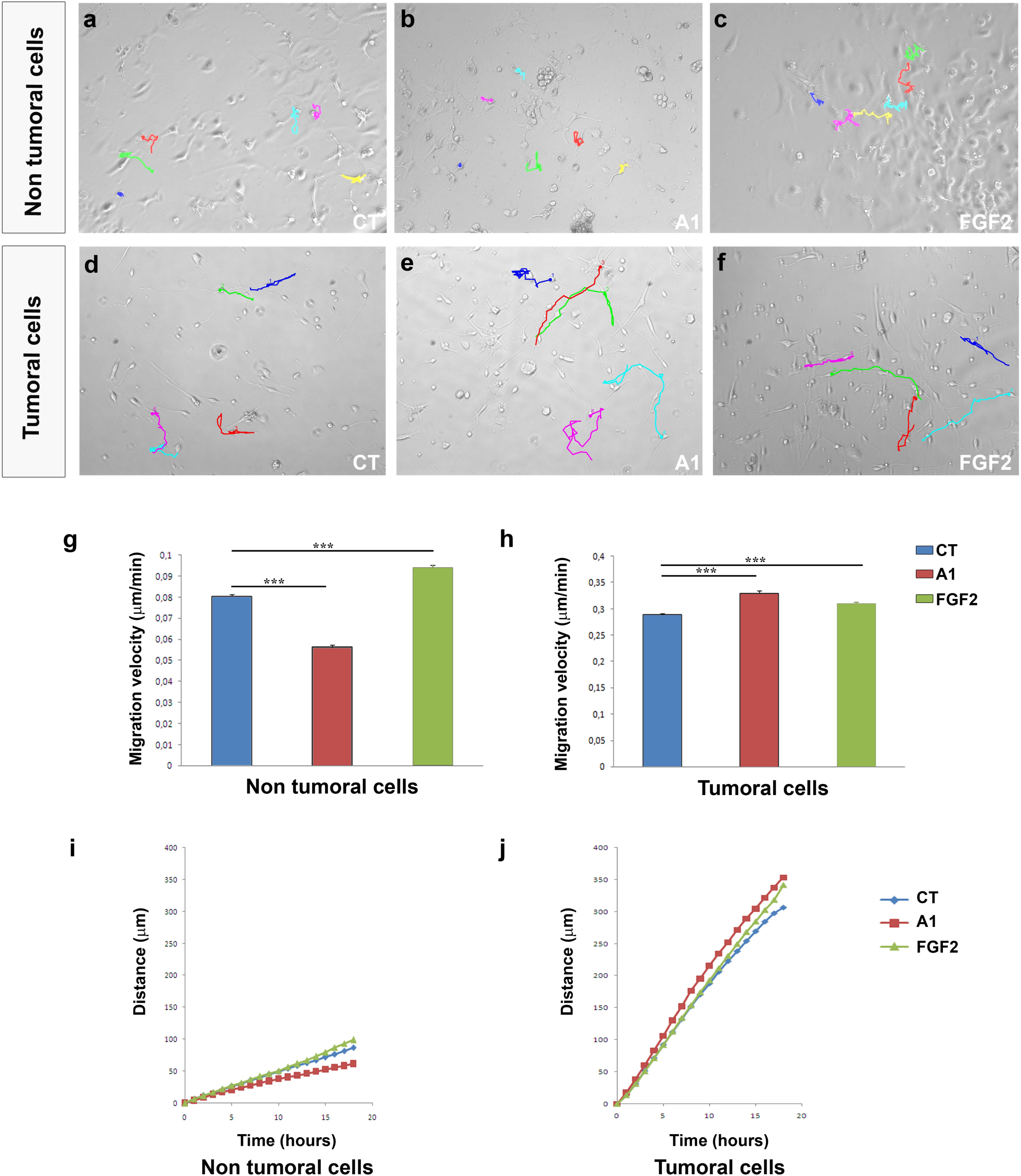
Video time lapse imaging of human OPC migration. (a-f) The manual tracking of non-tumor human OPCs (a-c) and tumor OPCs (d-f). Non-tumor cells migrate faster and further in the presence of FGF2 (g, i), while anosmin-1 has the opposite effect. Anosmin-1 and FGF2 both induce faster (j) and further migration (k) of tumor OPCs. The results were analyzed with a Student’s *t*-test: *P<0.05, **P<0.01, and ***P<0.001.

### The biological effects of FGF2 and anosmin-1 on OPC proliferation and differentiation

Rather than limiting our study to cell migration, we also studied how FGF2 and anosmin-1 affected OPC proliferation and differentiation. Both these factors had a constant effect on cell proliferation and while FGF2 significantly increased the number of newly-generated OPCs at each developmental stage analyzed and in young mature OPCs, anosmin-1 had no evident effect whatsoever on cell proliferation (Supplementary Fig.1a). Regarding OPC differentiation towards myelin-forming phenotypes, more heterogeneous results were obtained. In OPCs from embryonic (E16) and young adult (P60) stages, neither of these factors altered the rate of differentiation relative to the control conditions (Fig.6a-c, j-l, m). By contrast, FGF2 and anosmin-1 exerted opposite effects on P0 cells, while both these factors drove P15 OPCs towards a myelinating phenotype (Fig.6d-f, m).

**Figure 6:**
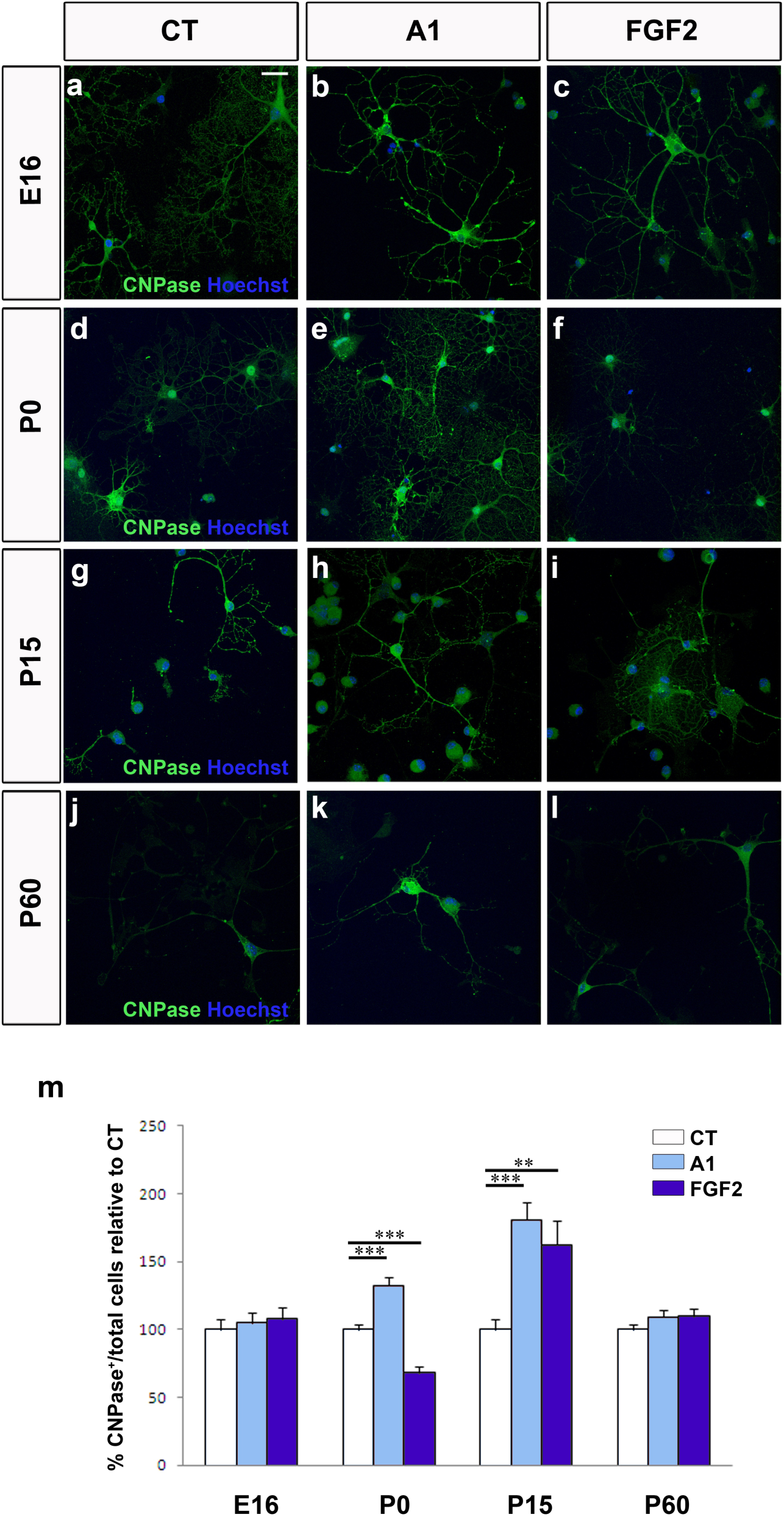
The effects of FGF2 and anosmin-1 on OPC differentiation. Images show mature oligodendrocytes from E16 (a-c), P0 (d-f), P15 (g-i) and P60 (j-l) mice labeled with CNPase (green) in control conditions and in the presence of FGF2 or anosmin-1. m) Histogram represents the proportion of CNPase^+^ cells relative to the controls at all the stages analyzed. Scale bar represents 25 µm for a-l and the results were analyzed with a Student’s *t*-test: *P<0.05, **P<0.01, and ***P<0.001.

### The effects of FGF2 and anosmin-1 on OPCs are FGFR-dependent

FGF2 and its receptors have been implicated in numerous processes, including cell proliferation, migration, differentiation and survival (for reviews see: [54-56]). Although we previously demonstrated that FGF2 and anosmin-1 exert their effects on OPC migration via FGFR1 [19,27,33,50,53], we analyzed the expression of the 3 FGFRs present in the oligodendroglial lineage [57,58] in our cultures of OPCs obtained from mice of different ages (E16, P0, P15 and P60). The FGFR1 was expressed by murine OPCs at every stage analyzed (Fig.7 a-l). Thus, to confirm that FGFR1 influences OPC migration, we repeated the chemotaxis assays on P0 (both anosmin-1 and FGF2 acting as chemoattractants) and P15 OPCs (anosmin-1 no effect; FGF2 chemoattractant), in the presence or absence of an FGFR signaling inhibitor (SU5402). This inhibitor abrogated both the chemoattraction and chemorepulsion provoked by FGF2 and/or anosmin-1, confirming FGFR1 as the FGF receptor involved in P0 and P15 OPC migration (Fig. 7m).

**Figure 7:**
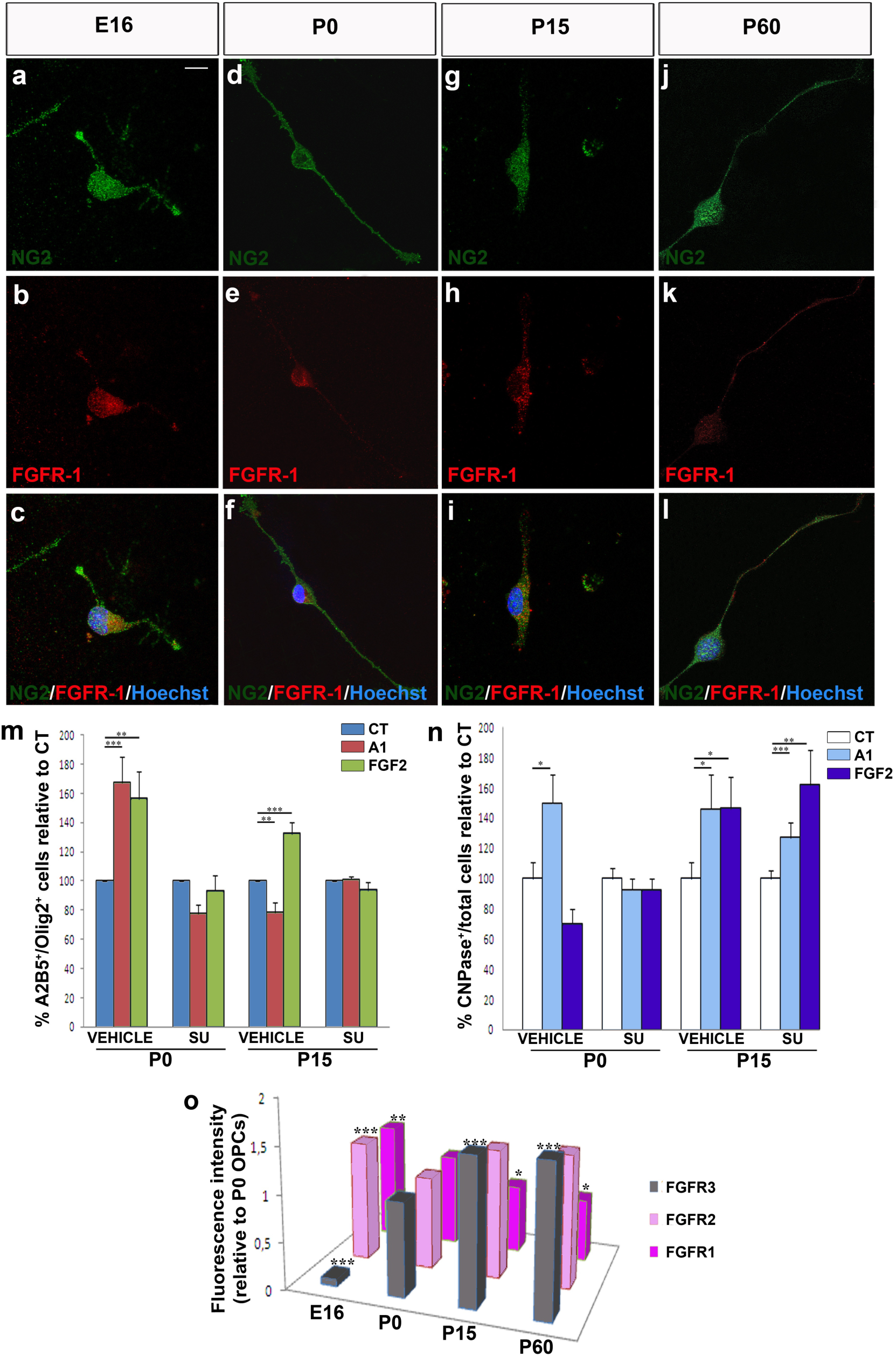
FGFR expression and its influence on OPCs. a-l) FGFR1 is expressed by OPCs at all the stages analyzed: E16 (a-c), P0 (d-f), P15 (g-i) and P60 (j-l). m) Histogram showing the results of chemotaxis at P0 and P15 in the presence of SU5402, which reverts the effects of anosmin-1 and FGF2. n) CNPase^+^ cells at P0 and P15 in cultures treated with SU5402 or the vehicle alone. No changes were observed in the presence of SU5402. o) FGFR1, R2 and R3 expression by OPCs isolated at E16, P0, P15 and P60. Scale bar represents 7.5µm for a-c, g-i, and 10 µm for d-f, j-l. The results were analyzed with a Student’s *t*-test: *P<0.05, **P<0.01, and ***P<0.001.

In terms of OPC differentiation, SU5402 abolished the effects of both molecules at P0, reducing the percentage of CNPase^+^ cells to control levels, although it failed to achieve the same effect on P15 OPCs exposed to either FGF2 or anosmin-1 (Fig.7n). As FGF2 binds to every FGFR but anosmin-1 binds exclusively to FGFR1 [52,53], it suggests that other FGFRs expressed by OPCs at this stage are likely to be involved in their differentiation [57-59]. Finally, when we analyzed the effect of SU5402 on OPC proliferation, we noted that the increase in proliferation observed in the presence of FGF2 was abolished with the inhibitor (Supplementary Fig. 1b).

Finally, we found a decrease in FGFR1 expression with age, which was most strongly expressed at the embryonic stage analyzed (Fig. 7o). This gradual decline mirrors the decrease in migration speed and chemotaxis in the presence of FGF2 and anosmin-1with age (Fig. 1). In parallel, FGFR2 expression was weakest at P0, with no remarkable changes at the other three stages studied (Fig. 7o). Finally, FGFR3 expression gradually increasing with age, from very weak expression at E16 to approximately 1.5-fold stronger expression at P15 and P60 than at P0 (Fig. 7o), consistent with previous reports [58].

## DISCUSSION

Given their relative abundance in the adult CNS, their ability to move towards lesion sites and their effectiveness in remyelination, there is ever increasing interest in adult OPCs. However, this is a highly heterogeneous cell population and in practical terms, the capacity of these cells to remyelinate lesions is limited [12,19,21,60-63]. Different developmental signaling cues, including growth factors, are over expressed locally in damaged areas, and they determine the survival, proliferation, migration and correct differentiation of endogenous adult OPCs [18,19,23,24,64] (for a review see [65,66]). Here we systematically studied heterochronic OPCs isolated from the mouse cerebral cortex at E16, P0, P15 and P60, as well as OPCs isolated from samples of adult human cerebral neocortex, avoiding the interference of the potential differences derived from the different developmental origins of the cells [2].

The heterogeneity in the responses of the OPCs also varied in function of the factor to which the cells were exposed. In response to FGF-2, OPCs responded in a more consistent manner when proliferation and migration were explored than when differentiation was studied. At each developmental stage studied, as well as in OPCs isolated from the cerebral cortex of adult mice and humans, FGF2 significantly enhanced cell proliferation (consistent with previous reports [55,67,68]) and certain migratory parameters, reflecting a motogenic and chemoattractive effect as reported previously [19,27,32,57,69]. It is interesting that the effect on human OPCs isolated from the safety-margins of tumors was, in absolute numbers, stronger than that on human adult OPCs isolated from non-tumor samples or even HOG cells (of human origin). It is also true that P15 OPCs had both a lower rate of proliferation and poorer mobility than the OPCs isolated from P0 and P60 mice. Our data on OPC differentiation towards myelin-forming phenotypes substantially differed with age. As such, while FGF2 interfered with OPC differentiation of P0 OPCs (as indicated elsewhere [59,70]), it favored the differentiation of P15 OPCs and it had a neutral effect on OPCs isolated from the adult cerebral cortex, independent of the species (similar results were obtained with rat P0 OPCs –data not shown- and adult human OPCs). NMR on murine OPCs also identified different metabolic patterns, and the less viscous cytoplasm of P15 OPCs compared to that of those isolated at P0 or P60 could explain the decreased in the migratory activity and proliferation OPCs at these stages [71-73]. This novel technique was previously used to analyze extracts of different cells types, including stem cells [74,75], although it was not applied to intact cells as occurred here.

In contrast to FGF2, anosmin-1 had no effect on OPC proliferation and it only potentiated OPC differentiation at early stages (P0, P15), having a neutral effect on differentiation at adult stages. However, the effects of anosmin-1 on OPC migration differed dramatically in function of both species and age, increasing the motility of P0 and tumor-related human OPCs (cells isolated from the safety-margins of tumors and also the HOG cell line), while it slowed down the migration of P15 and P60/adult OPCs, and that of OPCs from human non-tumor samples. These latter experiments confirmed earlier data and to some extent they explain the factors that condition OPC responses to anosmin-1 [27,32,33].

With respect to the different effects observed with human OPCs and other tumor cells, anosmin-1 seems to promote/attract the migration of tumor cells and immortalized cell lines ([52,76-79]; data herein). This might contribute to cancer cell dispersion and metastasis, while anosmin-1 can contribute to the immobilization of cells in both physiological conditions (data herein) and in non-tumoral diseases like MS [19]. Nevertheless, it was suggested elsewhere that the*Kal-1* gene would be a suppressor for other cancer types [78,80], which merits more systematic study. Likewise, the opposing or cooperative effects of FGF2 and anosmin-1 on tumor cells are still to be evaluated.

Undoubtedly, our study provides particularly interesting results in terms of the similarities and differences between OPCs isolated from the murine and human adult brain. The migratory properties of adult OPCs isolated from both species displayed parallel responses to FGF2 and anosmin-1, which is far from similar to the responses observed in early postnatal OPCs. Thus, our data demonstrate that OPC heterogeneity depends not only on the anatomical region from where they are isolated and on the way they respond to injury [21,81] but also, on the species, the stage in the life cycle and the functional system to which they pertain. This phenomenon should be taken into account in future studies that search for pharmacological agents to promote the motility of human OPCs isolated from different origins (the fetal or adult brain) in order to drive remyelination [66,82,83]. As such, and irrespective of their origin (murine or human), adult OPCs behave distinctly to OPCs isolated at earlier stages at least in part because their metabolic profiles are very different. In addition, their response to different cues, such as FGF2 and anosmin-1, is modified with age. Although not as systematic as our current work, earlier data also suggest that the effects triggered by growth factors and motogenic cues (PDGF, FGF2) are more constant than those driven by chemotropic molecules and/or cytokines (semaphorins, netrin, anosmin-1, CCLs [35,65,84,85]. Hence, these data together probably reflect a hierarchy among the different molecules that influence the biology of OPCs during development and in different species.

In conclusion, the data presented here help understand the response of endogenous OPCs to lesion in the adult CNS. As such, therapeutic approaches for neurodegenerative diseases should focus on the potentiation of the endogenous OPCs present in the adult brain, enhancing their spontaneous remyelinating capacity. Accordingly, it should also be borne in mind that the modulation of FGF family members might also improve myelin repair in diseases like MS.

## ABBREVIATIONS

CNS: Central nervous system
DMEM: Dulbecco’s Modified Eagle’s Medium
ECM: extracellular matrix
FBS: Fetal Bovine Serum
HRMAS: high-resolution magic angle spinning
MS: Multiple Sclerosis
OPC: Oligodendrocyte precursor cell

## AKNOWLEDGEMENTS

We thank Miss Iris Sánchez and Mr Rafael Lebrón for their technical support, as well as Dr José Ángel Rodríguez-Alfaro and Dr Javier Mazarío for their help with the imaging. This work was supported by grants from the Spanish Ministerio de Economía y Competitividad (MINECO: grant numbers SAF2009–07842, SAF2012-40023, RD07-0060-2007; and the Red Española de Esclerosis Múltiple: RD12-0032-12) and the ‘‘Fundación Eugenio Rodríguez Pascual’’ to FdC. ABA was hired under MINECO grants ADE10-0010 andSAF2012-40023. EMMR is the recipient of a predoctoral fellowship from the MINECO FPI program (BES-2010-042593 associated to SAF2009–07842). IM is contracted by SESCAM. FdC is a CSIC Staff Scientist.

