## Supplemental Figure 1 for "Age-linked heterogeneity among oligodendrocyte precursor cells in the cerebral cortex of mice and human"

**Supplementary Figure 1: Effects of FGF2 and anosmin-1 on OPC proliferation.** a) Quantification of the proliferating oligodendrocytes in the presence of FGF2 or anosmin-1 at the distinct ages studied. b) Graph showing how SU5402 influences the effect of FGF2 on the proliferation of OPCs. The results were analyzed using a Student´s *t*-test:*P<0.05, **P<0.01, and ***P<0.001.

**a**

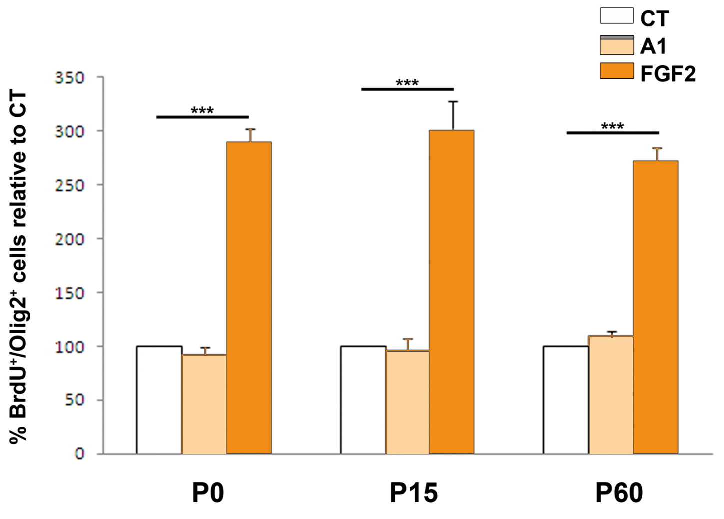


**b**

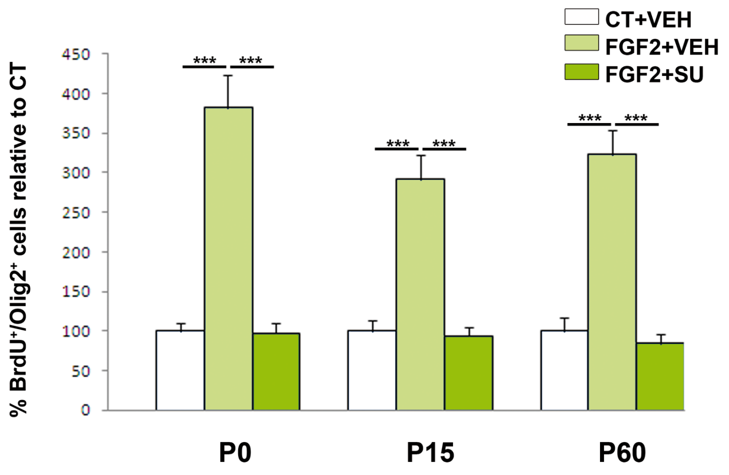
